# Complex eukaryotic-like actin regulation systems from Asgard archaea

**DOI:** 10.1101/768580

**Authors:** Caner Akıl, Linh T. Tran, Magali Orhant-Prioux, Yohendran Baskaran, Edward Manser, Laurent Blanchoin, Robert C. Robinson

## Abstract

Asgard archaea genomes contain potential eukaryotic-like genes that provide intriguing insight for the evolution of eukaryotes. The actin polymerization/depolymerization cycle is critical for providing force and structure for a variety of processes in eukaryotes, including membrane remodelling. Here, we identify actin filament severing, capping, annealing and bundling, and monomer sequestration activities by gelsolin proteins from Thorarchaeota (Thor), which complete a eukaryote-like actin depolymerization cycle. Thor gelsolins are comprised of one or two copies of the prototypical gelsolin domain and appear to be a record of an initial pre-eukaryotic gene duplication event, since eukaryotic gelsolins are generally comprised of three to six domains. X-ray crystal structure determination of these proteins in complex with mammalian actin revealed similar interactions to the first domain of human gelsolin. Asgard two-domain, but not one-domain, gelsolins contain calcium-binding sites, which is manifested in calcium-controlled activities. Expression of two-domain gelsolins in mammalian cells led to enhanced actin filament disassembly on ionomycin-triggered calcium release. This functional demonstration, at the cellular level, provides evidence for calcium-regulated actin cytoskeleton in Asgard archaea, and indicates that the calcium-regulated actin cytoskeleton predates eukaryotes. In eukaryotes, dynamic bundled filaments are responsible for shaping filopodia and microvilli. By correlation, the formation of the protrusions observed from Lokiarchaeota cell bodies may involve gelsolin-regulated actin structures.

---

Asgard archaea are some of the most fascinating organisms on the planet since they possess eukaryotic-like genes^1,2^, which encode for functional proteins^3^, providing evidence for the origins of the eukaryotic cell. In particular, Asgard genomes encode potential homologs for a regulated actin cytoskeleton^1,2^, which is critical for membrane remodeling in eukaryotes^4^, consistent with the membrane blebs, protrusions and vesicles observed in the first Asgard archaea to be isolated^5^. Asgard profilins support barbed-end actin filament elongation and inhibit spontaneous actin nucleation^3^. In addition, Asgard genomes also encode potential homologs for an ARP2/3 subunit and gelsolin domains^1–3^, which raises the question to what extent these archaea are able to regulate their actin dynamics beyond the control afforded by profilin^3^.

In eukaryotes, many actin-regulating activities, such as monomer sequestration and filament nucleation, elongation, annealing, bundling, capping and severing^6,7^, are elicited by the calcium-regulated multi-domain gelsolin family of proteins^8,9^. These proteins are predicted to have arisen from serial gene multiplication events^9,10^. Sequence and structure comparison of gelsolin domains indicates that two serial single domain gene multiplication events were followed by a third whole gene duplication to produce the 3 and 6 domain proteins, respectively^9–11^. The last common ancestor of eukaryotes likely possessed proteins of at least three gelsolin domains, since single or two domain proteins are not generally predicted from sequence databases. In addition, eukaryotes contain single domain ADF/cofilins and double cofilin domain twinfilins, some of which sever actin filaments and sequester actin monomers, and share a core domain topology with the gelsolin domain^12–16^.

## Results

We analyzed protein sequences from Asgard archaea genomes to search for proteins that contain gelsolin domains. We identified three gelsolin architectures from Thorarchaeota (Thor) sequence databases (Fig. 1a): a sequence comprising only of the prototypical gelsolin domain (ProGel); a single gelsolin domain followed by an unknown domain (1DGelX); and a two-domain gelsolin (2DGel). Similar architectures are found in, and are unique to, other Asgard archaea, including Lokiarchaeota. These mini gelsolins likely provide a record of the initial gene duplication in this superfamily of proteins (Fig. 1a). To date, the Asgard gelsolins are uncharacterized at the protein level. To further investigate the actin regulation in Asgard archaea, we tested the properties of Thor gelsolins.

**Fig. 1.**
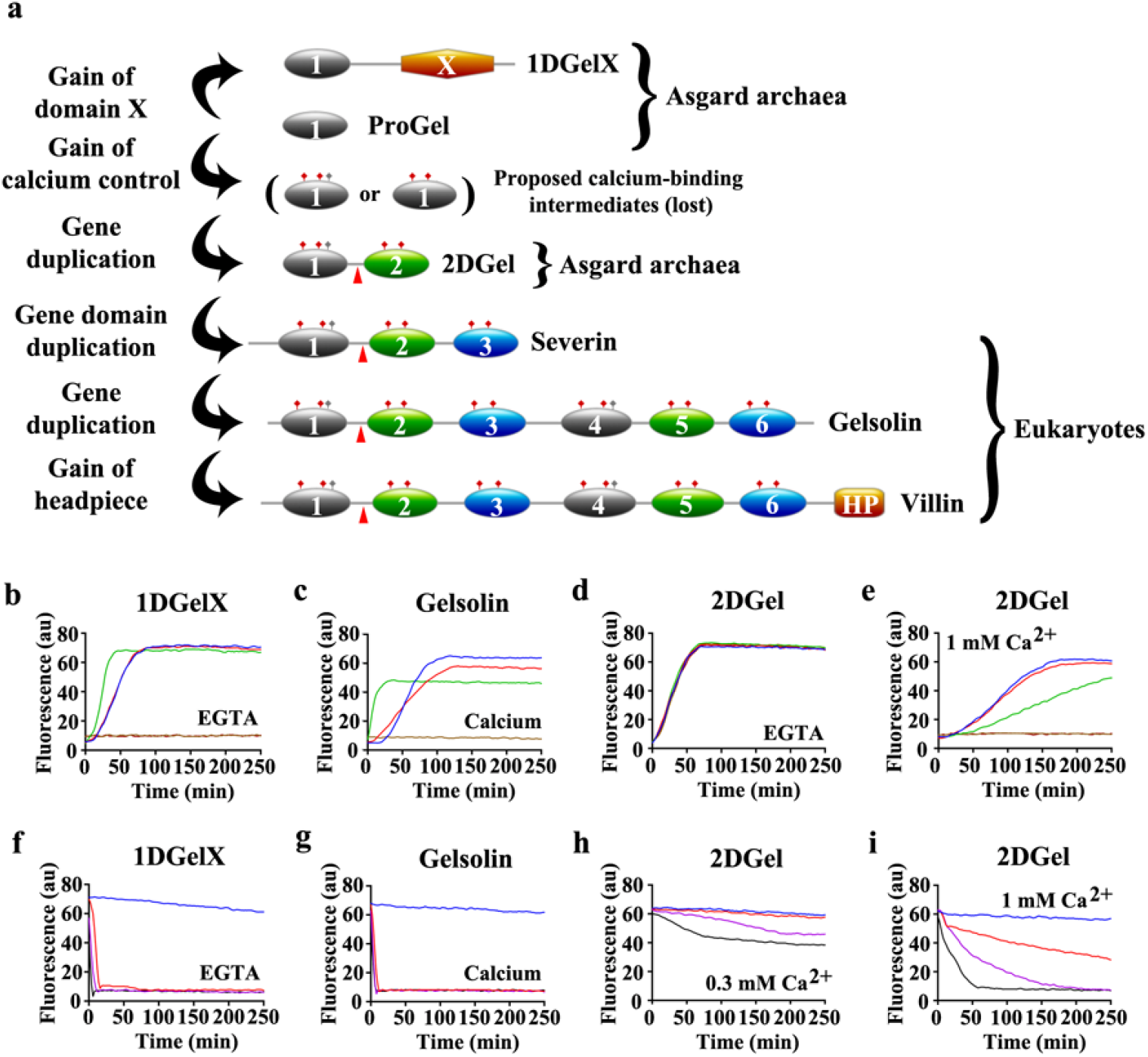
Thor gelsolins and actin regulation. **a**, Schematic representation of the three Thor gelsolin architectures. Ovals depict gelsolin domains. Ticks indicate potential calcium-binding residues and red triangles denote a central WH2-like motif. Since, Type II calcium-binding sites^25^ (red ticks) are found in both domains of 2DGel, a calcium-binding single-domain protein likely existed in evolution that is not found in the current sequence databases. This is indicated by the “proposed calcium-binding intermediates”. The Type I site^25^ (grey ticks) may have been present in this proposed calcium-binding intermediate, and later lost from domain two after the first gene duplication. Alternatively, the Type I site may have appeared in domain one after the first gene duplication. The architectures of typical eukaryotic gelsolin-like proteins are included for comparison. **b-e**, Pyrene-actin polymerization profiles of 2 μM actin (blue) supplemented with **b**, 1DGelX (1 mM EGTA), at 10 nM (red), 0.1 μM (green), or 2 μM (fawn) or 16 μM (dark brown), **c**, supplemented with 5 nM (red), 0.05 μM (green), 2 μM (fawn) human gelsolin (0.3 mM CaCl_2_), or supplemented with **d**, 2DGel (1 mM EGTA) or **e**, 2DGel (0.3 mM CaCl_2_) at the concentrations in (**b**). **f-i**, Actin depolymerization profiles of 2 μM actin (blue), supplemented by **f**, 1DGelX (1 mM EGTA), at 2 μM (red), 8 μM (lilac) or 32 μM (black), **g**, human gelsolin in 0.3 mM CaCl_2_, concentrations as in (**c**), **h**, 2DGel (1 mM EGTA) or **i**, 2DGel (0.3 mM CaCl_2_) at the concentrations in (**f**). Two other 2DGel orthologs, 2DGel2 and 2DGel3, which share 77-78% identity with 2DGel, showed additional filament nucleation activity and more potent severing activity. All three 2DGel proteins were less active at 0.3 mM than at 1 mM Ca^2+^, and inactive in 1 mM EGTA (Extended Data Fig. 1c-p).

We carried out *in vitro* biochemical experiments to demonstrate that Thor single-domain gelsolins are functional with eukaryotic actin in order to establish that these proteins are genuine gelsolin homologs. In pyrene-actin assays, 1DGelX and 2DGel showed robust activities. In an assembly assay, low concentrations of 1DGelX reduced the lag phase of actin polymerization consistent with filament nucleation, while 1:1 ratios inhibited polymerization (Fig. 1b), similar to calcium-activated gelsolin (Fig. 1c). The 1DGelX activity was similar in the presence of EGTA or Ca^2+^ (Extended Data Fig. 1a), indicating an absence of calcium control. By contrast, 2DGel showed no activity in the assembly assay in the presence of EGTA (Fig. 1d). However, in the presence of Ca^2+^ (1 mM), a low concentration of 2DGel (0.1 μM) significantly slowed actin polymerization indicating filament capping (Fig. 1e), and at a 1:1 ratio, no increase in pyrene fluorescence was observed, indicative of actin monomer sequestration. The sequestering and capping effects of 2DGel were reduced at intermediate Ca^2+^ levels (0.3 mM, Extended Data Fig. 1b).

In a pyrene-actin depolymerization assay, 1DGelX produced a rapid drop in fluorescence, indicative of robust severing (Fig. 1f), similar to calcium-activated gelsolin (Fig. 1g). 1DGelX’s severing activity was comparable in EGTA to Ca^2+^ (Extended Data Fig. 1c), revealing that this activity is Ca^2+^ independent. 2DGel in 0.3 mM Ca^2+^ showed partial loss in fluorescence implying incomplete depolymerization (Fig. 1h). At higher calcium levels (1 mM) the effects were more dramatic. 2DGel induced F-actin depolymerization with a fast, initial loss followed by a slower decline, consistent with filament severing followed by monomer sequestration (Fig. 1i). These effects were lost in EGTA (Extended Data Fig. 1d), implying that the calcium signalling range in Asgard archaea is likely to be higher than the micromolar range in some eukaryotes. ProGel had weak inhibitory effects on actin assembly and no effect on filament disassembly (Extended Data Fig. 2). These pyrene assay data reveal that 1DGelX and 2DGel display diverse gelsolin-like activities, including monomer sequestration and filament nucleation, capping and severing, with 2DGel showing calcium regulation.

Next, we applied TIRF microscopy to observe fluorescently-labelled actin filaments. Assembly of actin (1.5 μM) under high concentrations of ProGel (96 μM) confirmed an initial minor reduction in polymerization (0-4 min), but showed striking filament bundling and annealing at later time points under the assay’s molecular crowding conditions (0.25% methylcellulose, Fig. 2a, Supplementary Video 1). In this assay, increasing concentrations of 1DGelX (100 nM to 4 μM) produced a decreased number of filaments and some bundling at the highest concentration (4 μM, Extended Data Fig. 3b, Supplementary Video 2). 2DGel (24 μM) bundled filaments, with fewer structures in the presence of Ca^2+^ (Fig. 2a and Supplementary Video 3). The bundle morphologies appear strikingly similar to those observed for villin (Fig. 1a) in the TIRF microscopy actin assembly assay^17^.

**Figure 2.**
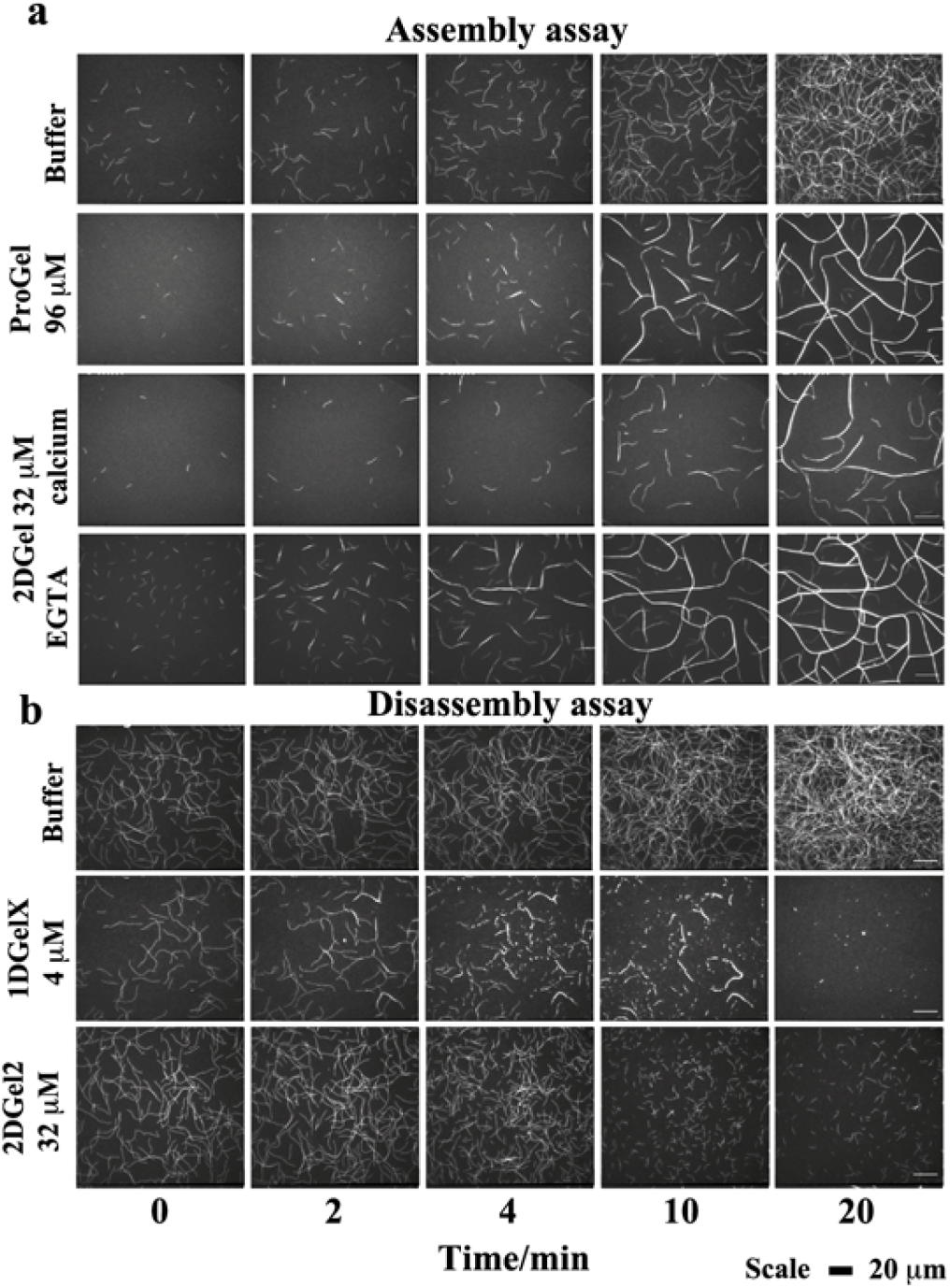
The regulation of actin assembly and disassembly by Thor followed by TIRF microscopy. Time course of the **a**, assembly and **b**, disassembly of 1.5 μM in the presence of various concentrations of Thor gelsolins. The scale bar represents 20 μM. Titrations of ProGel and 1DGelX in the assembly assay can be found in Extended Data Fig. 3, and comparison of 2DGel, 2DGel2 and 2DGel3 in the disassembly assay are found in Extended Data Fig. 4. Videos of the assembly/disassembly of Thor gelsolins are found in Supplementary Information Videos 1-5. ProGel and 1DGelX assays were carried out in 1 mM EGTA. The 2DGel assembly assays were in 0.3 mM CaCl_2_ or 1 mM EGTA, and the disassembly assay in 0.3 mM CaCl_2_.

Adding 1DGelX or 2DGel orthologs to preformed actin filaments resulted in severing by different modes. 1DGelX (4 μM) disassembled F-actin in a complex pattern (Fig. 2b and Supplementary Video 4). 1DGelX initially severed filaments into short fragments, which associated into short bundles, and the bundles were further severed to complete dissociation of actin structures. In 0.3 mM Ca^2+^, 2DGel (24 μM) bundled filaments, 2DGel2 (24 μM) severed single actin filaments with no bundling, and 2DGel3 (24 μM) initially severed single filaments, followed by the bundling and annealing of the fragments into larger structures (Fig. 2b, Extended Data Fig. 4 and Supplementary Video 5). These TIRF data reveal a complex regulation of the assembly and disassembly of actin filaments and bundles.

To compare the Asgard and eukaryotic gelsolin interactions with actin, we determined the X-ray structures of two types of Thor gelsolin/rActin complex. ProGel binds to actin at a site that overlaps with gelsolin domain 1 (G1)^18^, and with the cofilin family^19^ (Fig. 4a,d-f). Significantly, the protein is translated by one turn of the main helix relative to G1 and the calcium-binding sites are absent. The crystal structure of 2DGel bound to actin (Fig. 3b) revealed that domain 1 (D1) and the central WH2-like motif (LRRV) are similar to gelsolin (Fig. 3c), however, D2 is positioned differently^20^. ProGel and 2DGel contain the gelsolin DWG motif that is absent from the cofilin family (Extended Data Fig. 6a,b), implying that ProGel is a record of the ancestor of the gelsolin domain, despite its lack of calcium-binding activity. Next, we soaked the 2DGel crystals with 1 mM TbCl_3_ and examined the resultant anomalous electron density to confirm the cation-binding sites (Extended Data Fig. 6c)^21^. 2DGel binds four Ca^2+^, three sites in common with gelsolin (Extended Data Fig. 6d). We speculate that gene duplication to produce 2DGel resulted in tighter calcium control and extra functionality, due to multiplication of the calcium and actin-binding sites. Attempts to crystallize 1DGelX failed. The 1DGelX gelsolin domain shares 47% identity with ProGel and is likely to similarly bind to actin (Fig. 3a). Domain X is predicted to form a coiled-coil structure with no homology to known actin-binding proteins (Extended Data Fig. 5c). Given 1DGelXs robust actin-filament severing activity (Fig. 1g), domain X is likely to interact with filaments, however we were unable to produce the domain X protein to verify this point.

**Figure 3.**
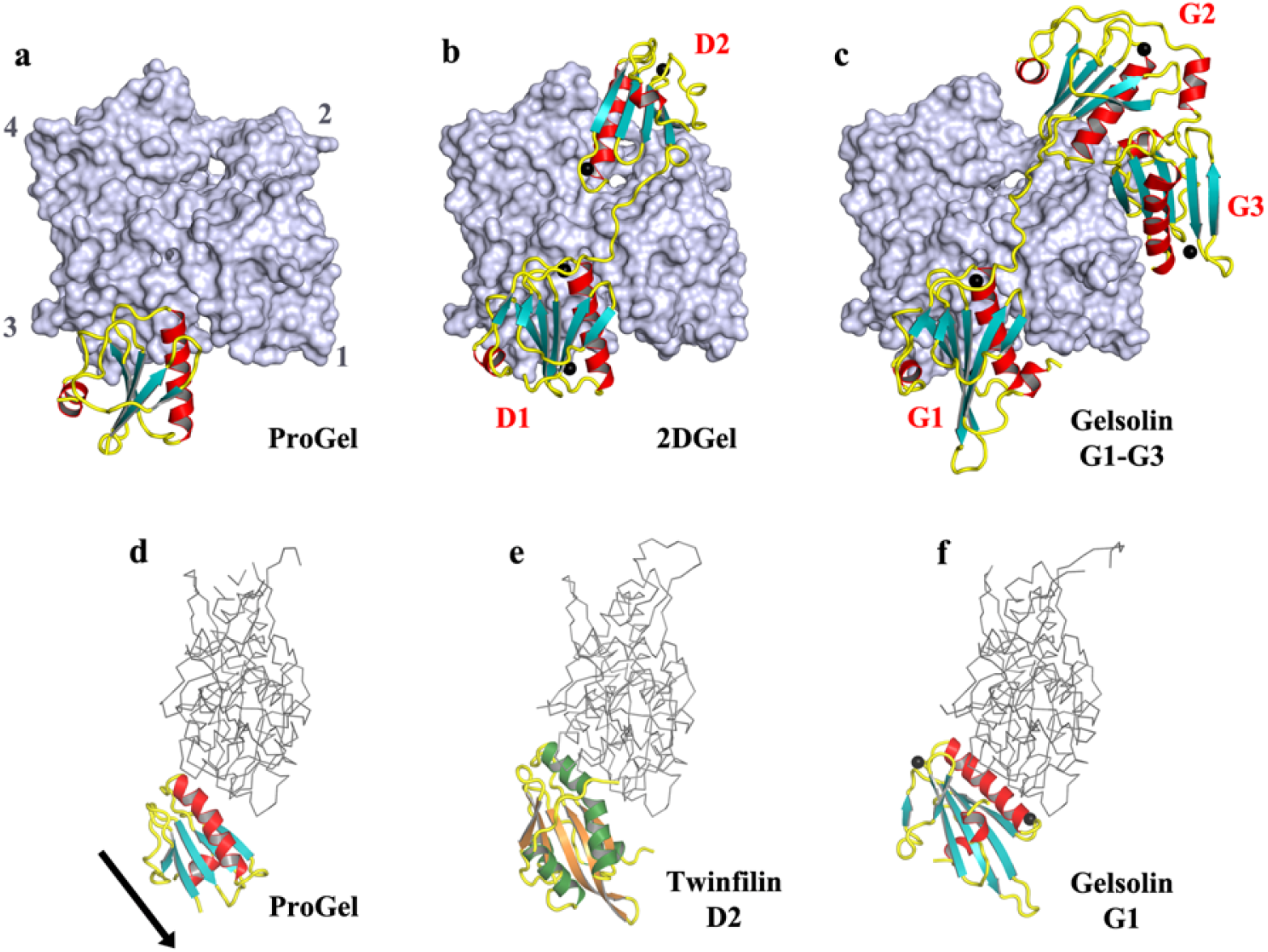
The structures of ProGel and 2DGel in complex with rActin. **a**, The ProGel/rActin complex. rActin is shown as a surface and ProGel is in schematic representation. **b**, The structure of the 2DGel/rActin complex. The four calcium ions associated with 2DGel are shown as black spheres. The crystal structure of 2DGel3/rActin complex is found in Extended Data Fig. 5a. **c**, The structure of the first three domains of human gelsolin in complex with rActin for comparison (PDB code 1EQY). **d-f**, Side views of **d**, ProGel, **e**, twinfilin domain 2, a cofilin-family member (PDB code 3DAW), and **f**, gelsolin domain G in complex with actin, with actin shown as a trace. Similar representations for 2DGel and 2DGel3 are found in Extended Data Fig. 5b. The arrow indicates the displacement of ProGel relative to G1. Data collection and refinement statistics are found in Extended Data Table 1. Further analyses of the structures are found in Extended Data Fig. 5,6.

**Figure 4.**
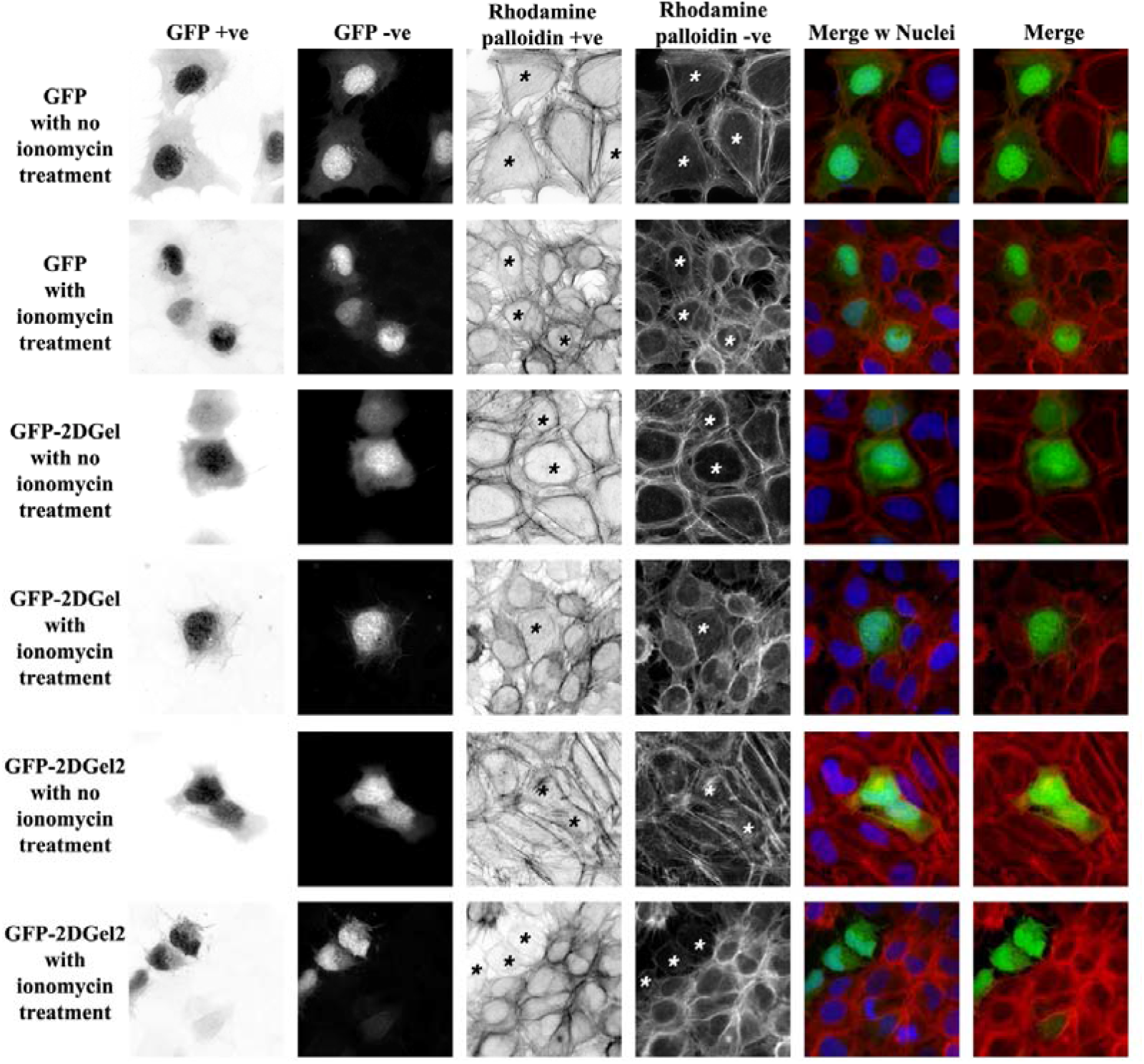
Calcium signalling to ectopically expressed 2DGel proteins in human U2OS cells followed by fluorescence imaging. Cell expressing GFP, GFP-2DGel or GFP-2DGel2 are indicated by signal the GFP channel, columns 1 and 2, and highlighted with asterisks. Actin filaments and larger structures are observed in the rhodamine-phalloidin channel, column 3 and 4. “+ve” refers to normal and “-ve” to the reversed image. Merged images of the GFP channel (green) and the rhodamine-phalloidin channel (red) are shown with (blue) or without nuclei staining in the final two columns. Different cells were imaged immediately before or 10 minutes after treatment with ionomycin to release calcium. Statistics are found in Extended Data Fig. 7.

Finally, we tested whether 2DGel could respond to calcium signalling in a cellular context. GFP fusion constructs of 2DGel, and its ortholog 2DGel2, were transfected into human U2OS cells and the cells were treated with rhodamine-phalloidin, which stains actin filaments. GFP positive cells, showed typical cell morphologies, actin cytoskeleton arrangements, and GFP localization for all constructs (Fig. 4). Treatment of the cells with ionomycin (10 μM, 10 min), to release calcium from intracellular stores, led to a contraction of all cells. In the GFP-2DGel and GFP-2DGel2 expressing cells, but not GFP-expressing cells, a dramatic loss in rhodamine-phalloidin signal was observed (Fig. 4 and Extended Data Fig. 7). This indicates that calcium signalling to 2DGel and 2DGel2 resulted in the loss of F-actin structures, demonstrating that these proteins are functional in the eukaryotic cellular environment.

## Discussion

These data indicate that Asgard mini gelsolins represent a record of the early gene duplication events in the gelsolin family of proteins (Fig. 1a) and demonstrate calcium signalling as a control mechanism for the Asgard cytoskeleton. Asgard mini gelsolins possess equivalent actin-regulating properties to their larger multi-domain eukaryotic gelsolin counterparts, and to proteins that possess different architectures, such as capping protein and fascin (Fig. 5). We speculate that the emergence of diverse actin regulators in eukaryotes allowed for greater and more distinct control of actin dynamics, enabling its incorporation into an expanded number of cellular processes22. However, the fundamental actin architectures, actin filaments and filament bundles, likely existed in pre-eukaryotic organisms. The profilin and gelsolin controlled actin polymerization/depolymerization cycle in Asgard archaea, and the assembly/disassembly of higher order structures, indicate a complex actin cytoskeleton (Fig. 5). We now have an emerging picture of a highly regulated Asgard eukaryotic-like polymerization/depolymerization cycle that is executed by a limited number of protein folds, yet has many of the characteristics of the actin dynamics in eukaryotes (Fig. 5), where a large part of the functional output involves eliciting membrane perturbations, similar to those observed in Lokiarchaeota5. Furthermore, this analysis of Thor gelsolins highlights the evolution of the gelsolin and cofilin families of proteins. The core structure of ProGel is similar to the cofilin and gelsolin domains (Extended Data Fig. 6e). This may indicate that present day eukaryotic cofilins and gelsolins evolved from a protein that had similar properties to ProGel. The acquisition of sequences outside of the core ProGel domain appears to be a critical factor in extending the range of functions. Eukaryotic cofilin has additional terminal helices whereas eukaryotic gelsolins have duplications of the core domain. Comparison of the activities of ProGel with 1DGelX and 2DGel indicates how addition of extra domains to the core gelsolin domain leads to altered activities. Thus, ProGel may represent a record of an initial cofilin/gelsolin protein that originally emerged as a relatively simple actin regulator, which through gaining C-terminal domains, or expansion of the core, added additional functionalities.

**Figure 5.**
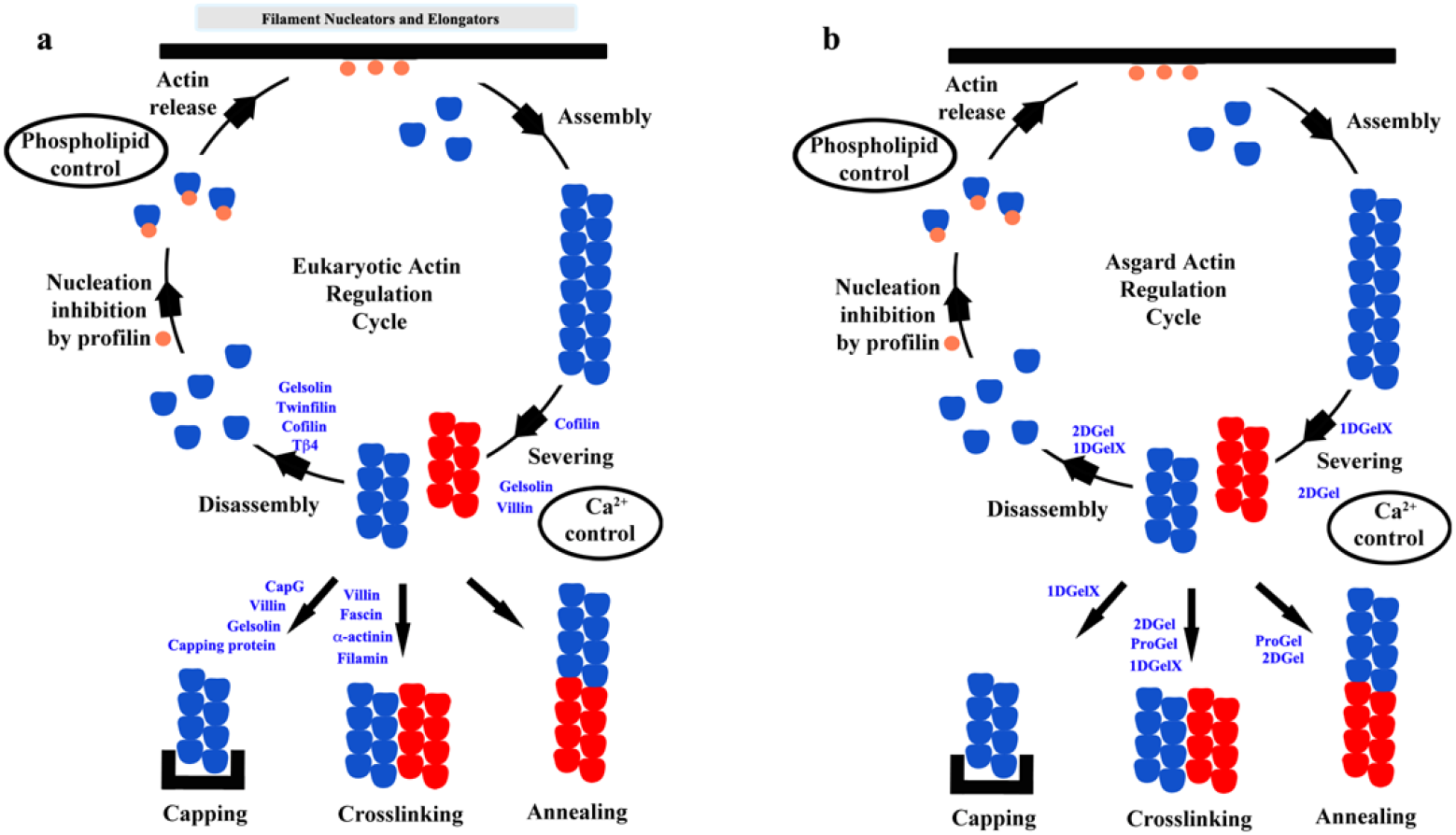
Comparison of the actin polymerization cycle between a, eukaryotes and b, Asgard archaea. Eukaryotes have specialized proteins, formed from a variety of folds, that have superseded and added further sophistication to the profilin^3^ and gelsolin domain-based actin regulation determined for Asgard archaea. For instance, CapG is a capping protein, gelsolin severs and caps filaments, whist villin also bundles filaments, all under calcium control, whilst, capping protein or fascin do not require calcium for activity.

ProGel, 1DGelX, 2DGel and Asgard profilins3 possess sequences and structures that are unique to Asgard archaea, with orthologous sequences also found in Heimdallarchaeota, Lokiarchaeota and Odinarchaeota genomes, however these proteins are not predicted from eukaryotic genomes. Furthermore, similar gelsolin and profilin sequences are present in the genome of the first Asgard archaea to be isolated and sequenced5. Hence, these genes are not eukaryotic contaminants from metagenomics analyses. The presence of functional gelsolin-like proteins in Asgard archaea has several possible scenarios with regards to the emergence of the domains of life, which has been debated in relation to different phylogenetic analyses3,5,23,24. In the three-domain hypothesis, eukaryotes and archaea form separate clades. In this case, there are at least two possible explanations for the existence of Asgard gelsolins that are capable of interacting with eukaryotic actin. Firstly, gelsolins, profilins and actins may have been present in last common ancestor of archaea and eukaryotes, but later lost from most branches of archaea, with the exception of Asgard archaea. Such wide-spread loss of these genes in Archaea would seem unlikely given the prevalence of these proteins throughout eukaryotes, indicating the advantage of these genes for survival. Secondly, the genes may have been passed by horizontal gene transfer to stem Asgard archaea from a stem eukaryote(s) or vice versa. We specify stem organisms, since all Asgard phyla contain ProGel and 1DGelX, and eukaryotes generally contain three domain gelsolins, requiring that that the last common ancestor of each domain would contain these genes. Given the abundance of eukaryotic-like protein sequences in Asgard archaea genomes, this would require a sizeable transfer, or transfers, of genetic material. Alternatively, in the two-domain hypothesis, eukaryotes emerge from within the archaea domain, sharing a common ancestor with Asgard archaea. In this scenario, which involves the least caveats, the gelsolin, profilin and actin proteins arose in a common ancestor of eukaryotes and Asgard archaea. Regardless of the phylogeny, the primitive gelsolins analysed here represent a record of a nascent actin cytoskeleton regulation machinery that was likely a prerequisite for eukaryogenesis.

## Acknowledgements

We thank A*STAR for support and William Burkholder for reagents. We thank the experimental facility and the technical services provided by: The Synchrotron Radiation Protein Crystallography Facility of the National Core Facility Program for Biotechnology, Ministry of Science and Technology and the National Synchrotron Radiation Research Center, a national user facility supported by the Ministry of Science and Technology, Taiwan, ROC; and by the Australian Synchrotron, part of ANSTO. We thank Professors Jian-Ren Shen and Yuichiro Takahashi for use of reagents and access to equipment.

## Author Contributions

C.A. and R.C.R. conceived experiments. C.A., L.T.T., M.O.-P., Y.B. and R.C.R. performed experiments. M.O.-P. and L.B. conceived and performed TIRF microscopy experiments. Y.B. and E.M. conceived and performed cell imaging experiments. All authors analysed data. All authors contributed to writing the paper.

## Author Information

The authors declare no competing financial interests. Correspondence and requests for materials should be addressed to R.C.R.

## Methods

### Protein expression and purification

Gene sequences were synthesized and codon optimized for *E. coli* (GenScript) and placed in the pSY5 vector^3,26^. Proteins were expressed as described previously^3^. Asgard gelsolin cell pellets (10 g) were resuspended in binding buffer 50 ml (20 mM HEPES, pH 7.5, 500 mM NaCl, 20 mM imidazole, 1 mM TCEP and 1 mM EGTA), supplemented with benzonase (2 μl of 10000 U/μl, Merck), Protease Inhibitor Cocktail (Set III, EDTA-Free, Calbiochem) and Triton X-100 (0.01%). Cell lysis, Ni-NTA affinity and size-exclusion chromatography (16/60 Superdex 75 PG, GE Healthcare) in gel filtration buffer (20 mM HEPES, pH 7.5, 150 mM NaCl, 1 mM TCEP and 1mM EGTA) were performed as described previously^3^. Asgard gelsolin fractions were pooled and concentrated (10000 MWCO Vivaspin concentrator, Vivascience). Human gelsolin was purified as described previously^26^. Freshly prepared rActin was purified including a final gel filtration step, and was labelled with pyrene as described previously^27^. For TIRF studies, actin was labelled on lysines by incubating actin filaments with Alexa-488 succinimidyl ester (Molecular Probes)^28^. Human profilin was expressed in BL21 DE3 pLys S cells and purified as reported^29^. Attempts to express Thor actin and constructs comprising individual 1DGelX gelsolin and X domains failed.

### Pyrene-actin assays

Pyrene-actin assembly and disassembly assays were performed with 2 μM rabbit skeletal-muscle G-actin (10% pyrene labelled). Gelsolin polymerization and depolymerization assays were performed in the presence of Ca^2+^ or EGTA. In the pyrene-actin assembly assays, G-actin in buffer A (2 mM Tris.HCl, pH 7.4, 0.2 mM ATP, 0.5 mM DTT, 0.3 mM or 1 mM CaCl_2_, 1 mM Na azide) was mixed and incubated with a 20-fold dilution of 20X Mg-exchange buffer (1 mM MgCl_2_, 4 mM EGTA) for 2 min in order to pre-exchange the calcium ion for magnesium. Subsequently, actin polymerization was initiated by the addition 10 μl of 10X actin polymerization buffer (500 mM KCl, 10 mM MgCl_2_, 100 mM imidazole-HCl, pH 7.5) supplemented by the appropriate concentrations of EGTA or CaCl_2_ in a total volume of 100 μl.

Pyrene-actin disassembly assays were carried out in presence of calcium (0.3 mM or 1 mM Ca^2+^) or EGTA (1 mM or 2 mM). For pyrene-actin disassembly assays, 2 μM G-actin (10% pyrene labelled) was mixed with buffer A. Actin polymerization was initiated by the addition of 10 μl of 10X KMEI or KMI (KMEI without EGTA) buffer. The polymerization was monitored around 2 h at room temperature in 96-well, black, flat-bottomed plates in total volume of 90 μl. Then, 10 μl of different concentrations of gelsolin proteins in buffer A were added to the preformed F-actin in the appropriate concentrations of CaCl_2_ or EGTA. All reactions were performed in 96-well, black, flat-bottomed plates (Corning, Nunc). The fluorescence intensities were monitored at wavelength 407 nm after excitation at 365 nm with a Safire^2^ fluorimeter (Tecan). Seeding-pyrene assays were performed as described^3^.

### TIRF assays

20 × 20 mm^2^ coverslips and cover glasses (Agar Scientific) were extensively cleaned, oxidized with oxygen plasma (5 min at 80 W, Harrick Plasma, Ithaca, NY) and incubated with 1 mg/ml of silane-PEG, MW 5K (Creative PEG Works) overnight. Actin assembly was initiated in polymerization chambers of 20 × 20 mm^2^ × 4.5 μm height or in a PDMS chamber with a reaction volume of 30 μL, by addition of the actin polymerization mix (2.6 mM ATP, 10 mM DTT, 1 mM EGTA, 50 mM KCl, 5 mM MgCl_2_, 10 mM HEPES, pH 7.5, 3 mg/mL glucose, 20 μg/mL catalase, 100 μg/mL glucose oxidase, 0.2% w/v BSA and 0.25 % w/v methylcellulose) containing actin monomers (1.5 μM, 20% Alexa488-labeled).

The polymerization chambers were constructed using a double-sided tape (70 mm height) between a glass coverslip and slide coated with silane-PEG. The PDMS chambers were prepared using the published protocol^30^. To observe the effect on actin polymerization, Thor gelsolins were added at the desired concentration directly in the polymerization mix. To show effect on F-Actin, the PDMS chamber was used to reconstitute actin network during 5 to 10 min, then the gelsolin protein was added carefully and mixed. The controls were carried out using gelsolin storage buffer (20 mM HEPES, pH 7.5, 150 mM NaCl, 1 mM TCEP and 1 mM EGTA) without gelsolin.

### Structure determination, model building and refinement

Native X-ray diffraction datasets from single crystals of 2DGel/rActin and 2DGel3/rActin on a RAYONIX MX-300 HS CCD detector on beamline TPS 05A (NSRRC, Taiwan, ROC) controlled by BLU-ICE (version 5.1) at λ□=□1.0 Å. Data were indexed, scaled, and merged in HKL2000 (version 715)^31^ (Extended Data Table 1). A native data X-ray diffraction dataset from a crystal of the ProGel/rActin complex was collected on beamline MX2 (Australian Synchrotron) on a Eiger16M detector at λ□=□1.0 Å. Data were indexed, scaled, and merged in XDS (Version November 2016)^32^ and ccp4-7.0 CCP4-7.0 AIMLESS (version 0.5.29) (Extended Data Table 1). Terbium anomalous diffraction data for 2DGel/rActin were collected and merged in the range 20.0-1.7 Å (Rmerge 0.056, Rpim 0.056, anomalous completeness 99.4%, redundancy 5.5) using the same protocols as for the native data set.

Molecular replacement using the ProGel/rActin and 2DGel/rActin; datasets using the native actin (PDB code 3HBT)^27^ was carried out in the PHENIX suite (version 1.13-2998)^33^ Phaser. The model for 2DGel/rActin was extended in AutoBuild^33^, whereas the gelsolin chains for ProGel/rActin were built by hand. All manual adjustments to the models and refinement were carried out in Coot (version 0.8.9 EL)^34^. Final refinement for ProGel/rActin was carried out in CCP4-7.0 Refmac5^35^. The final model for 2DGel/rActin was used as the molecular replacement search model for 2DGel3/rActin. All final models were verified for good stereochemistry in PHENIX suite (version 1.13-2998)^33^ MolProbity^36^ (Supplementary Table 1). The final ProGel/rActin models (2 in the asymmetric unit) consist of ProGel residues 1-88 missing the final 4 residues (chain B and D) and rActin 5-42 and 51-374 (chain A) and rActin 5-41 and 51-374 (chain C). The actins are each associated with ATP, latrunculin B and a magnesium ion. Electron density for the 24 residues of the ProGel models. The final 2DGel/rActin model consists of 2DGel residues 2-197 (missing the final residue), associated with 5 calcium ions, and rActin 5-41 and 49-372. The final 2DGel3/rActin model consists of 2DGel3 residues 2-198, associated with 5 calcium ions, and rActin 6-39 and 51-372.

### Sequence analyses

Protein domain homolog identification was carried in BLAST^37^ using reported Asgard sequences^1,2^. Domain architectures of proteins were created in Prosite MyDomains^38^.

### Cell culture, transfection and intracellular calcium induction

U2OS (human osteosarcoma) cells were cultured in high glucose Dulbecco’s modified Eagle’s (DME) media with 4500 mg/L glucose, supplemented with 10% Fetal Bovine Serum (FBS) (Hyclone). Cells were grown at 37°C in an incubator filled with 5% CO_2_ and 99% humidity. U2OS cells were seeded onto 22 × 22 mm glass coverslips in 35 mm culture dishes and grown to sub-confluence. Each dish was then transfected with 0.5 μg of GFP control or fusion gelsolin construct plasmid DNA using the Mirus TransIT LT1 transfection reagent according to the manufacturers protocol, incubated at 37°C and allowed to express for 24 hours. Cells were observed to ensure at least >50% of cells were expressing GFP, appeared healthy and of comparable density. The media was then changed to reduced serum DME (1% FBS, 1.8 mM calcium) with or without 10 μM ionomycin (a calcium ionophore) and incubated at 37°C for 10 min. Cells were then fixed with 3.7% formaldehyde in PBS at room temperature for 20 min.

### Statistics and Reproducibility

All biochemical experiments were repeated 3 times with similar results.

---

**Extended Data Figure 1.**
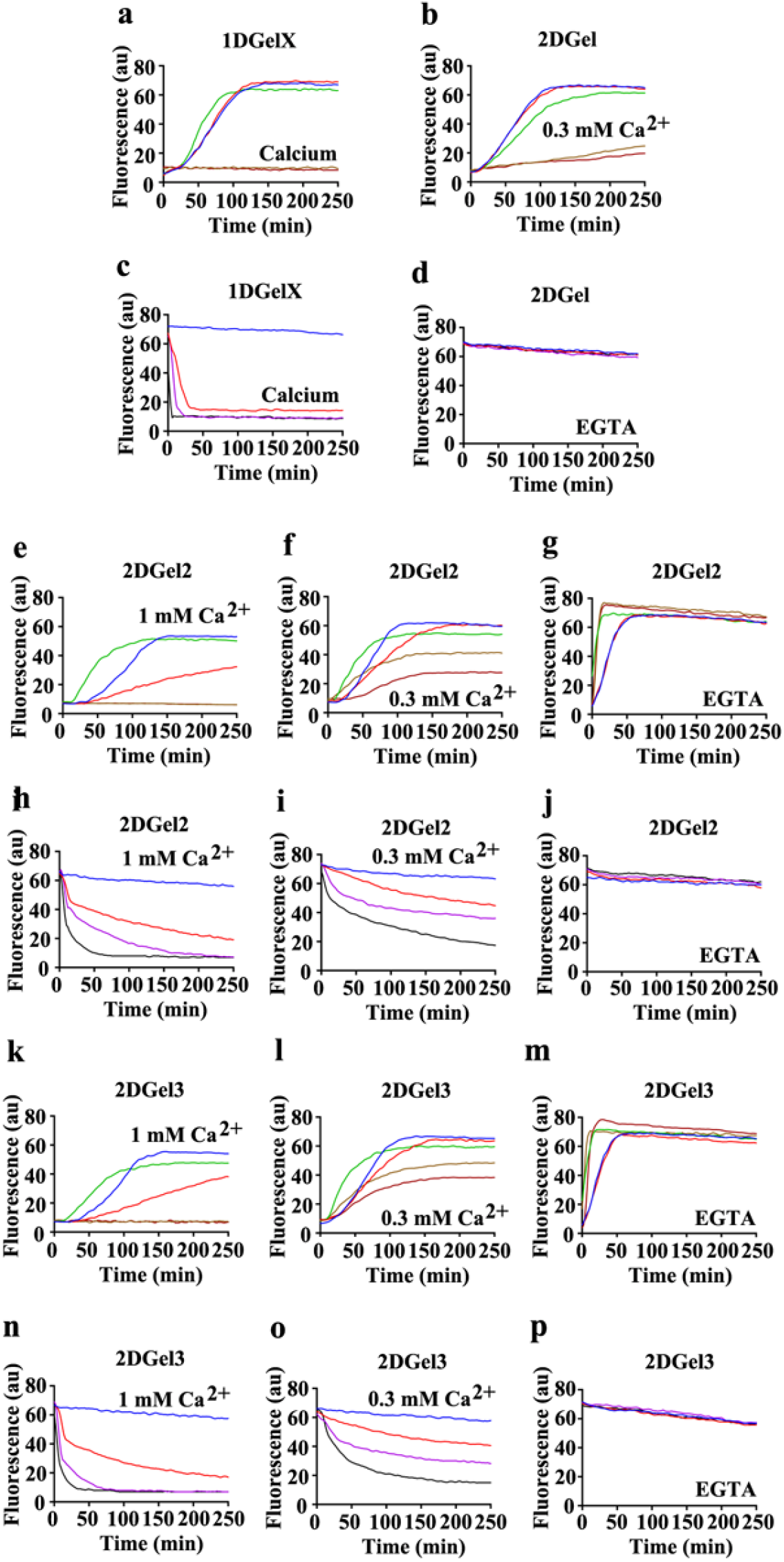
Actin regulation by 2DGel2 and 2DGel3 and control experiments for Fig. 1. Concentrations and solution conditions are a detailed in Fig. 1.

**Extended Data Figure 2.**
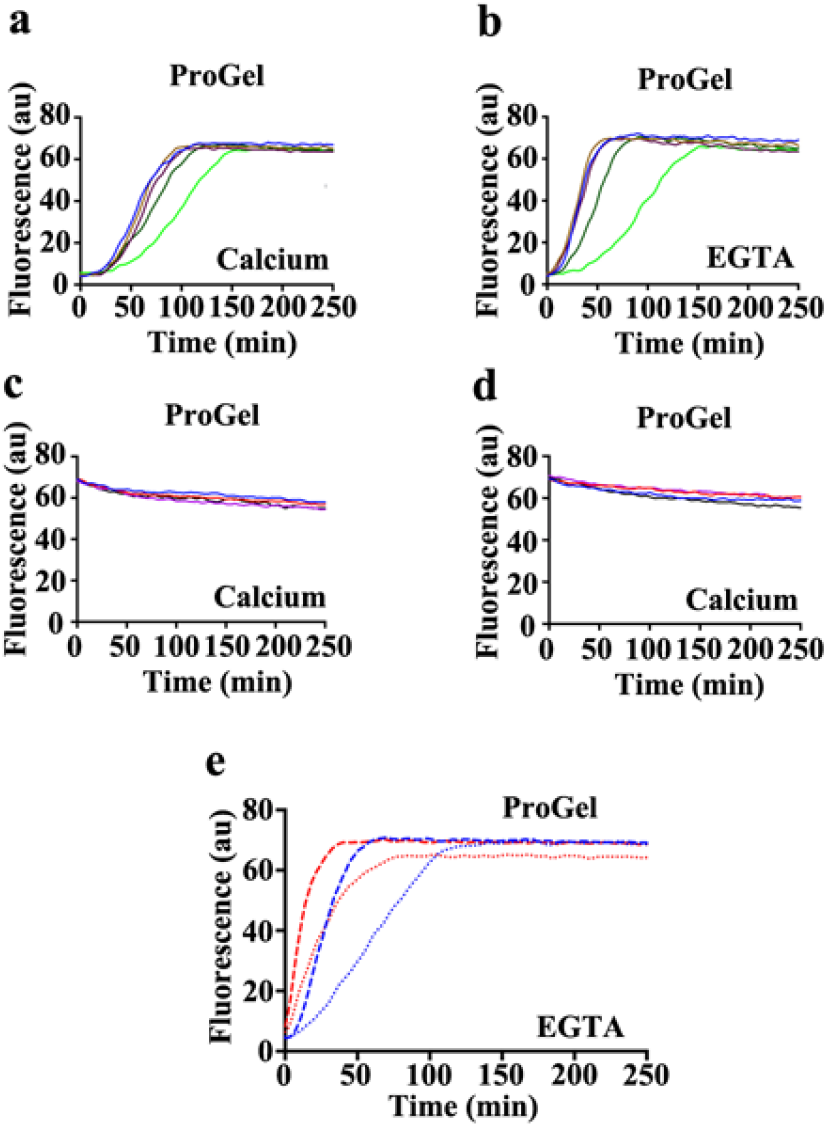
Actin regulation by ProGel. **a,b**, Pyrene-actin polymerization profiles of 2 μM actin (blue) supplemented with 2 μM (fawn), 8 μM (lilac), 32 μM (dark green) or 128 μM (light green) ProGel in **a**, 1 mM EGTA or **b**, 0.3 mM Ca^2+^. The high concentrations of ProGel produced a delay in the polymerization-induced fluorescence, although the final levels attained were similar, under both calcium and EGTA conditions. **c,d**, Pyrene-actin depolymerization profiles of 2 μM actin (blue) supplemented with 10 nM (red), 0.1 μM (green), 2 μM (fawn) or 16 μM (dark brown) ProGel in **c**, 1 mM EGTA or **d**, 0.3 mM CaCl_2_. No effect was observed in this pyrene actin disassembly assay. **e**, Pyrene-actin polymerization profiles of 2 μM actin (dashes) or 2 μM actin with 128 μM ProGel (dots) in the presence (red) or absence (blue) of actin filament seeds. The delay in fluorescence increase (**a,b**) could be overcome by adding actin filament seeds. These data suggest that ProGel may have partial profilin-like properties, as observed for the Asgard profilins^3^, in supporting filament elongation, but suppressing spontaneous actin nucleation, or may have monomer sequestering or filament capping properties, and these properties are not under calcium control. However, the effects on rabbit actin were very weak and Thor actin is required to confirm whether these are genuine properties of ProGel.

**Extended Data Figure 3.**
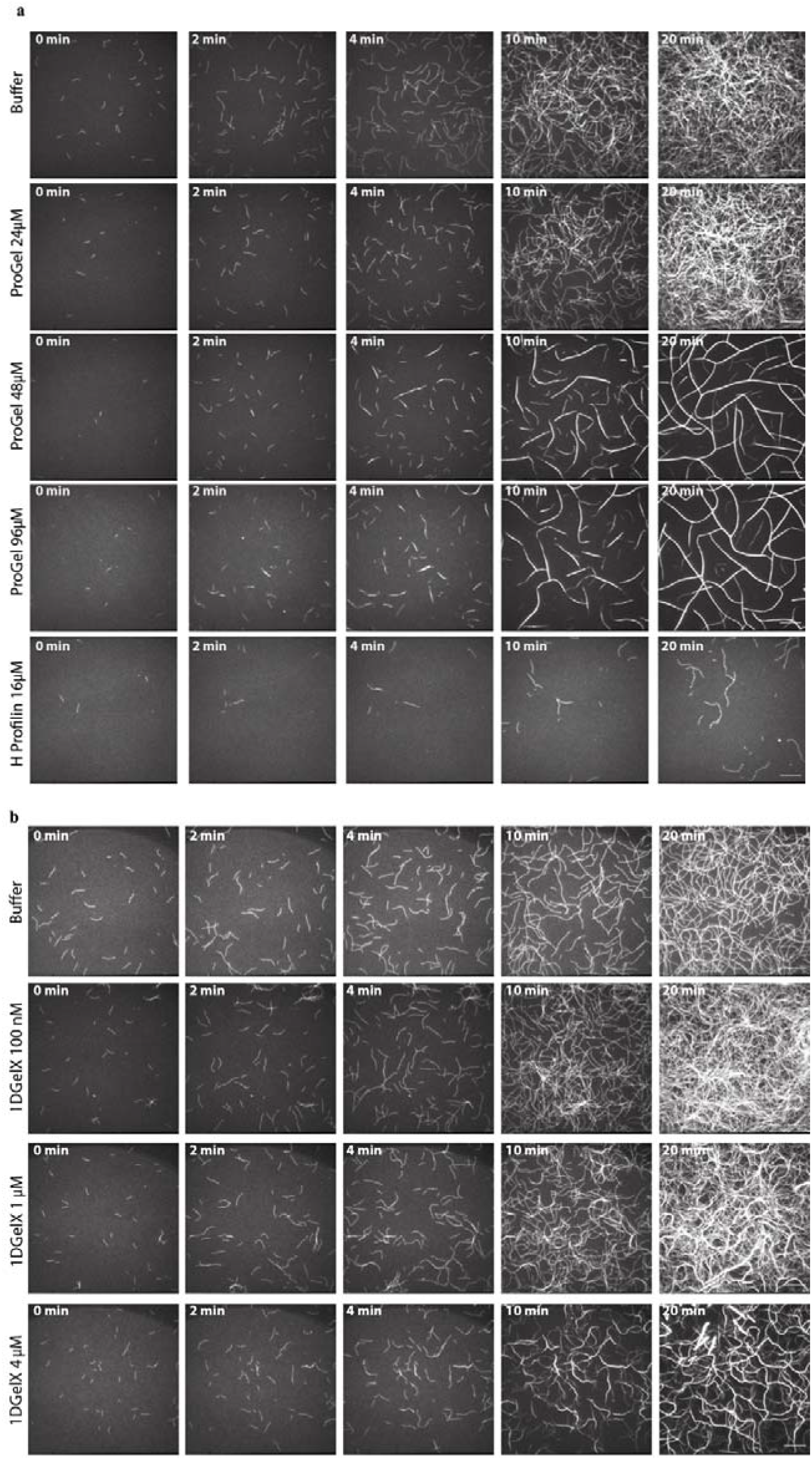
Effects of ProGel and 1DGelX on actin polymerization. Time course of polymerization of 1.5 μM actin in the presence of **a**, various concentrations of ProGel or human profilin (16 μM), **b**, various concentrations of 1DGelX. The figures were generated from Supplementary Videos 1 and 2, respectively. The scale bar represents 20 μM.

**Extended Data Figure 4.**
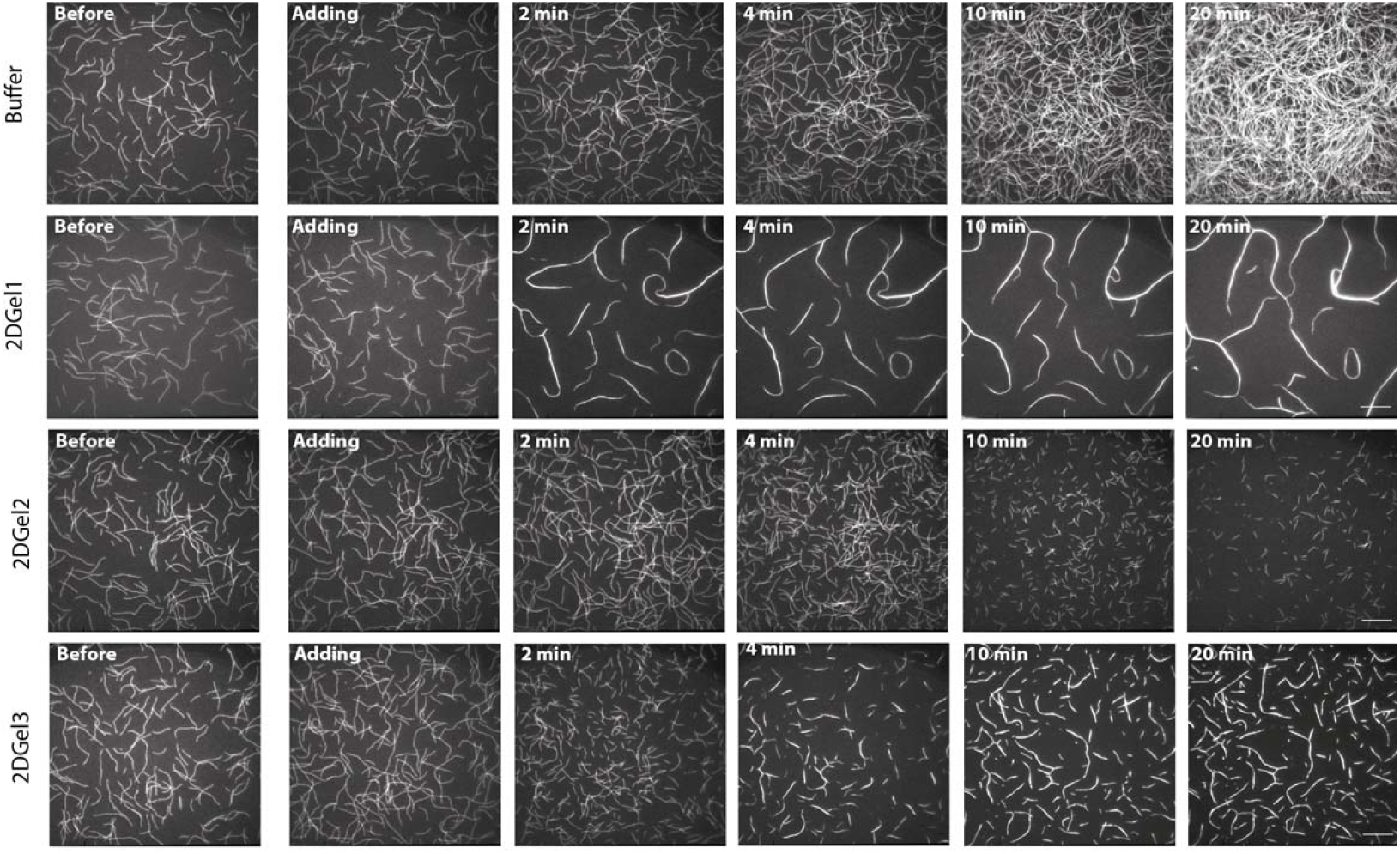
Effects of 2DGel, 2DGel2 and 2DGel3 on actin depolymerization. Time course of depolymerization of 1.5 μM actin in the presence of 24 μM 2DGel proteins in 0.3 mM CaCl_2_. The figure was generated from Supplementary Video 5. The scale bar represents 20 μM.

**Extended Data Figure 5.**
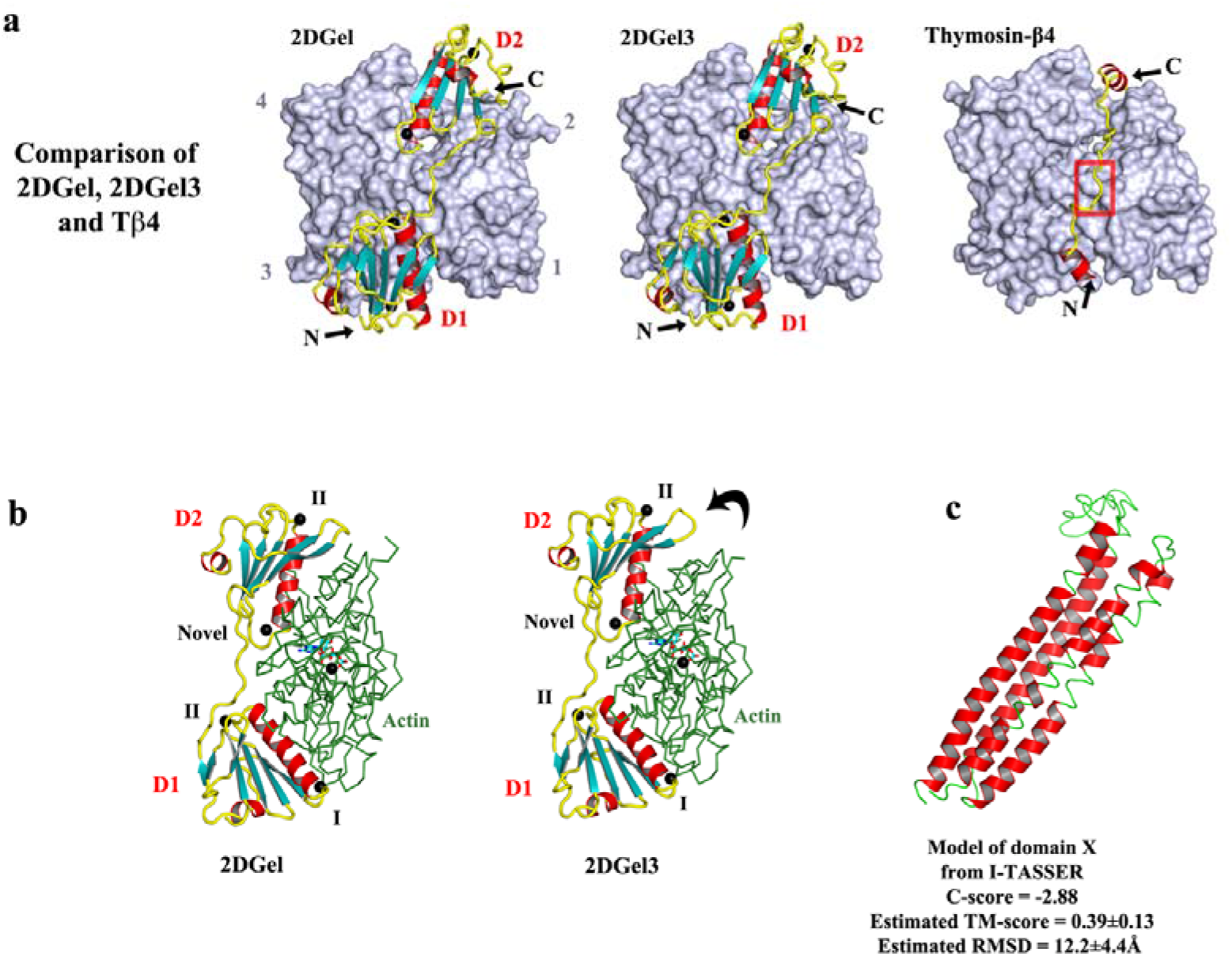
Comparison of the structures of 2DGel and 2DGel3 in complex with actin. **a**, The 2DGel, 2DGel3 (a 2DGel ortholog) and Tβ4 (PDB code 4PL7) structures in complex with actin. Actin is shown as surfaces and binding partners are shown in schematic representation. The red box indicates the position of the “LKKT” WH2-like motif on the thymosin-β4 structure. D1, D2, N, C and numbers indicate domain 1, domain 2, N-terminus, C-terminus, and the subdomains of actin, respectively. **b**, Side on views of the 2DGel/rActin complexes. I, II and novel refer to Type I, Type II or the novel calcium-binding sites, respectively. These Type I and Type II calcium-binding sites are conserved in human gelsolin. D2 from 2DGel packs more closely to the surface of actin than D2 form 2DGel3 (indicated by the arrow), possibly providing a structural basis for the difference in activities of these proteins (Extended Data Figure 4). **c**, Model of the structure of domain X from 1DGelX generated by I-TASSER, with statistics^39^.

**Extended Data Figure 6.**
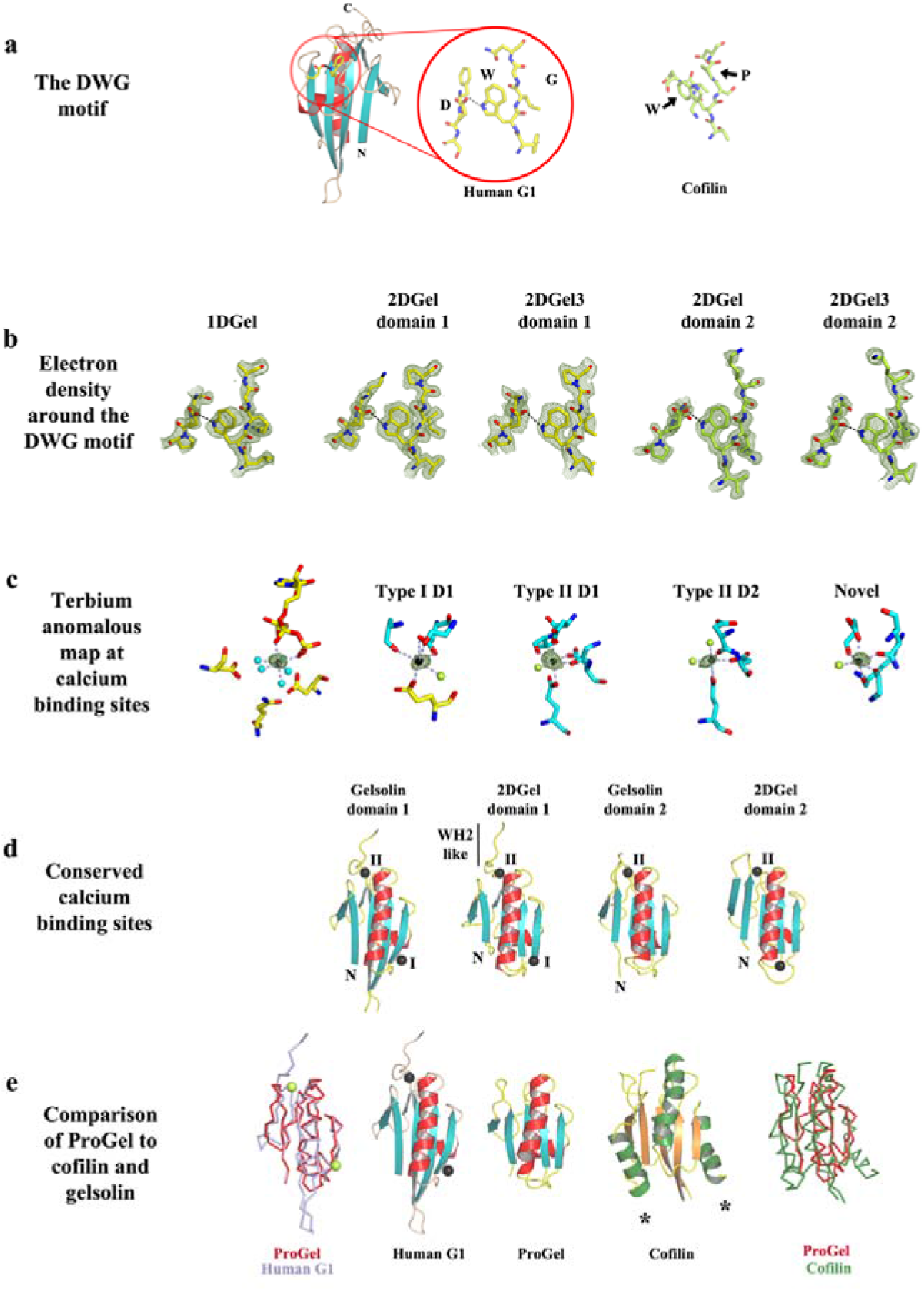
The DWG motif and calcium-binding sites. **a**, The DWG motif. The location of the DWG motif, in which the aspartic acid forms a hydrogen bond to the tryptophan which sterically occludes residues larger than glycine. This is present in all domains of human gelsolin (PDB code 3FFN) but absent from the cofilin fold, such as mouse twinfilin D2 (PDB code 3DAW). **b**, The OMIT map electron density (contour level 1 σ) is shown around the DWG motifs from the Thor gelsolins. Middle panel. **c**, The terbium anomalous difference map (contour level 6-8 σ) showing density at each of the metal ion binding sites in the 2DGel/actin complex. Actin residues are shown in yellow, 2DGel in cyan, Tb^3+^ as black spheres and waters as light green or cyan spheres. **d**, The 3 conserved calcium-binding sites in 2DGel and 2DGel3, shared with the first two domains of human gelsolin. **e**, Structural comparisons and superimpositions of ProGel with human gelsolin G1 (PDB code 3FFN) and cofilin (PDB code 4KEE). Cofilin likely arose from a ProGel-like protein, since they share a conserved structural core. Calcium ions are shown as lime or black spheres. Asterisks indicate additional helices in the cofilin fold relative to ProGel.

**Extended Data Figure 7.**
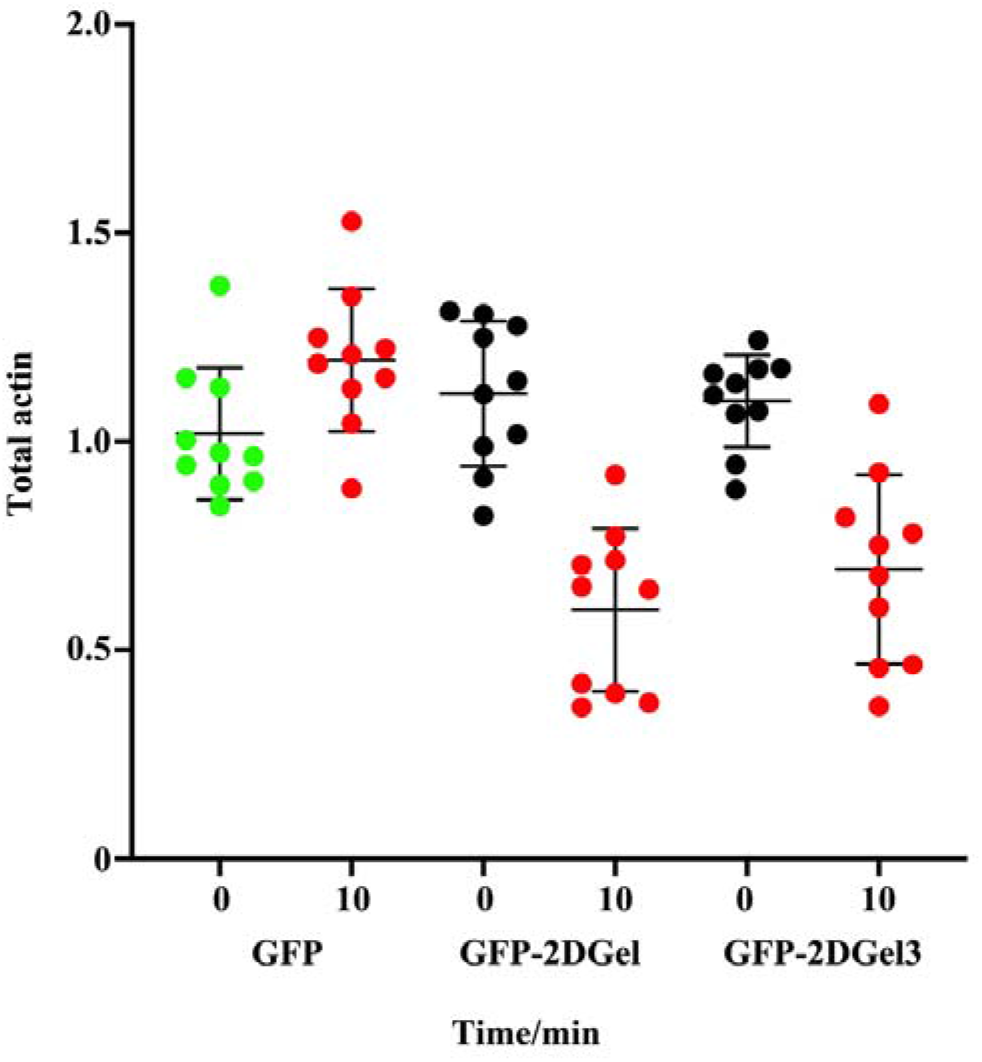
Quantification of rhodamine-phalloidin fluorescence before and 10 minutes after treatment with ionomycin. 12-bit monochrome images of actin fluorescence intensity, of typical cells, were quantified as a ratio for GFP and adjacent non-GFP expressing cells, in 3 separate experiments.

**Extended Data Table 1.** X-ray data collection and refinement statistics.

|  | ProGel/rActin | 2DGel/rActin | 2DGel3/rActin |
| --- | --- | --- | --- |
| **Data collection** |  |  |  |
| Crystal | P2<sub>1</sub>2<sub>1</sub>2<sub>1</sub> | P2<sub>1</sub> | P2<sub>1</sub> |
| *a*, *b*, *c* (Å) | 57.4, 108.6, 167.5 | 52.9, 101.3, 55.8 | 49.9, 98.6, 62.5 |
| $\alpha$, $\beta$, $\gamma$ (°) | 90.0, 90.0, 90.0 | 90.0, 96.6, 90.0 | 90.0, 94.0, 90.0 |
| Wavelength (Å) | 1.0 | 1.0 | 1.0 |
| Resolution (Å)<sup>a</sup> | 91.1-2.03 (2.08-2.03) | 20.0-1.71 (1.73-1.70) | 31.2-2.35 (2.44-2.35) |
| *R*<sub>merge</sub> | 11.5 (79.0) | 10.0 (92.3) | 13.5 (58.4) |
| *R*<sub>meas</sub> | 12.5 (85.6) | 10.8 (108.0) | 14.4 (64.3) |
| *R*<sub>pim</sub> | 4.8 (32.7) | 4.2 (45.5) | 5.5 (26.3) |
| *I*/$\sigma$(*I*) | 8.6 (1.8) | 20.1 (1.8) | 17.3 (2.3) |
| *CC*<sub>1/2</sub> | 0.998 (0.785) | 0.926 (0.632) | 0.946 (0.862) |
| Completeness (%) | 99.5 (93.0) | 98.0 (92.5) | 92.4 (81.4) |
| Redundancy | 6.7 (6.6) | 6.6 (4.5) | 6.8 (5.5) |
| **Refinement** |  |  |  |
| Resolution (Å) | 91.1-2.03 (2.08-2.03) | 20-1.71 (1.74-1.70) | 31.2-2.35 (2.48-2.35) |
| No. reflections | 64952 (4484) | 58175 (1354) | 23279 (2562) |
| *R*<sub>work</sub> / *R*<sub>free</sub> | 21.9/24.3 (33.0/31.8) | 16.9/20.3 (22.4/26.4) | 20.2/23.8 (29.6/34.2) |
| No. atoms* |  |  |  |
| Protein | 5709 (A)/1428 (G) | 2943 (A)/1620 (G) | 2823 (A)/1615 (G) |
| Ligand/ion | 116 (LatB, ATP)/2 (Mg<sup>2+</sup>) | 31 (ATP)/7 (Ca<sup>2+</sup>) | 31 (ATP)/6 (Ca<sup>2+</sup>) |
| Water | 535 | 557 | 124 |
| *B* factors |  |  |  |
| Protein | 27.3/27.8 | 33.5/37.0 | 19.6/24.9 |
| Ligand/ion | 22.3/12.2 | 26.0/18.9 | 15.2/11.3 |
| Water | 37.5 | 36.9 | 33.6 |
| r.m.s deviations |  |  |  |
| Bond lengths (Å) | 0.009 | 0.007 | 0.009 |
| Bond angles (°) | 1.43 | 0.88 | 1.05 |
| Ramachandran Plot |  |  |  |
| Favoured (%) | 97 | 97 | 98 |
| Outliers (%) | 0 | 0 | 0 |

\* ProGel/rActin has 2 complexes in the asymmetric unit.

## Supplementary Videos

**Supplementary Information Video 1** | Time course of polymerization of 1.5 μM actin supplemented by buffer, ProGel (24 μM), ProGel (48 μM) or ProGel (96 μM). ProGel (24 μM) shows slightly reduced polymerization relative to the control. ProGel at 48 μM and 96 μM shows bundling and annealing. All movies are sped up 200 times.

**Supplementary Information Video 2** | Time course of the polymerization of 1.5 μM actin in the presence of buffer, 1DGelX (100 nM), 1DGelX (1 μM), or 1DGelX (4 μM). 1DGelX at 100 nM shows filament nucleation. 1DGelX at 1 μM and 4 μM show increasing levels of bundling.

**Supplementary Information Video 3** | Time course of the polymerization of 1.5 μM actin in the presence of buffer, 2DGel (32 μM) and 1 mM EGTA, or 2DGel (32 μM) and 1 mM CaCl_2_. 2DGel in the 1 mM CaCl_2_ shows reduced polymerization and bundling in comparison to the presence of 1 mM EGTA.

**Supplementary Information Video 4** | Disassembly of polymerized actin (1.5 μM) produced by 1DGelX (4 μM) in comparison to the buffer control. 1DGelX shows single filament severing, bundling followed by bundle severing.

**Supplementary Information Video 5** | Polymerized actin (1.5 μM) in the presence of 0.3 mM CaCl_2_ supplemented by buffer, 2DGel (32 μM), 2DGel2 (32 μM) or 2DGel3 (32 μM). 2DGel shows bundling, 2DGel2 shows single filament severing, and 2DGel3 shows single filament severing followed by bundling.

